# Stimulating a NTS to LC circuit elicits anxiety-like behavior and state-dependent autonomic output

**DOI:** 10.64898/2026.09.30.755810

**Authors:** A.G. Butler, J.K. Bassi, A.A. Connelly, M. R. Melo, A.M. Allen, S.J. McDougall

## Abstract

Understanding how the brain integrates internal bodily signals to shape behavior is essential for characterizing the neural basis of interoception. Here, we investigate whether the nucleus of the solitary tract (NTS), a brainstem hub for viscerosensory input, communicates with the locus coeruleus (LC) to influence anxiety-like behavior. We utilized viral-induced optogenetics combined with electrocardiography and diaphragmatic electromyography to monitor heart rate and respiratory frequency. Prior to behavioral testing we photoactivated transduced NTS neurons either at their somata, or at their axonal terminals in the LC. Anxiety-related behaviors were assessed using the elevated plus maze (EPM) and novelty-suppressed feeding test (NSFT). Stimulation of the NTS somata, or their terminals in the LC, increased aspects of anxiety-like behaviors in the EPM and NSFT, including freezing time during the EPM and latency to eat in the center of the NSFT. Importantly, only NTS somal activation suppressed post-NSFT feeding. This suggests that the anxiety-like behaviors observed from activating NTS projections to the LC are not secondary to activating feeding circuits. Additionally, activation of NTS terminals in the LC decreased heart rate only following food deprivation. These findings reveal that the NTS to LC circuit enhances anxiety-like behavior and modulates cardiovascular dynamics state dependently, thereby identifying a likely brainstem pathway through which viscerosensory signals influence interoception.

**Significance Statement:** Interoception is a fundamental brain function that allows sensory signals from internal organs to shape perception and behavior. Despite its importance, the circuit mechanisms that distribute viscerosensory signals across the brain remain poorly understood. Here, we show that a projection from the nucleus of the solitary tract (NTS) to the locus coeruleus (LC) increases aspects of anxiety-like behavior and alters heartrate in a state-dependent manner. These findings causally identify a novel pathway that enables visceral signals to access a major neuromodulatory center and influence behavior and cardiovascular dynamics, thereby providing a foundational framework for understanding how interoceptive signals and circuits shape both anxiety and automatic output.

## Introduction

Interoception is fundamental to survival, continuously shaping perception, autonomic control, and behavior (Craig, 2002; Critchley & Garfinkel, 2017). Interoception depends, in part, on a continuous flow of sensory feedback from vagal viscerosensory neurons, with each heartbeat, breath, and gut contraction providing visceral information that allows the brain to monitor and regulate internal states. Following processing by brainstem circuits, viscerosensory signals are relayed to higher-order networks involved in emotion and cognition (Critchley & Harrison, 2013). Dysfunction in these networks contributes to a range of neuropsychiatric disorders and maladaptive behavioral states (Khalsa et al., 2018; Bonaz et al., 2021). Therefore, the identification of the specific neural circuits that contribute to interoceptive function is crucial for understanding how internal physiological states shape cognition and behavior.

The nucleus of the solitary tract (NTS) is a primary brainstem nucleus that receives viscerosensory inputs, processes the information and then relays it to higher order nuclei. Whilst traditionally described as an autonomic center, the NTS is both anatomically and functionally heterogeneous, contributing to a broad range of behaviors in addition to classical homeostatic functions (Holt, 2022). For example, selectively modulating NTS neuron activity with chemogenetic or optogenetic approaches regulates feeding (Aklan et al., 2020; D’Agostino et al., 2016; Roman et al., 2017; Sayar-Atasoy et al., 2023) and affective behavior (D’Agostino et al., 2016; Chen et al., 2020; Roman et al., 2017). These findings support the idea that interoceptive signals processed and relayed upstream via the NTS can modulate both affect and behavior to support homeostatic functions. However, the different nuclei that receive and integrate input from the NTS to elicit interoceptive outcomes remain incompletely characterized.

The noradrenergic locus coeruleus (LC) is a neuromodulatory center that may be important for linking interoceptive signals to cognition. It receives dense input from both autonomic and cognitive regions and can exert widespread influence through broad projections across the neuroaxis (Poe et al., 2020). The LC also modulates a range of cognitive and affective processes, including arousal, attention, learning, memory, behavioral flexibility and anxiety-like behavior (Poe et al., 2020; Suárez-Pereira et al., 2022; Boyle et al., 2024; Bouras et al., 2023; Bari et al., 2020). In addition, LC firing responses to food are influenced by fed state (Sciolino et al., 2022) and LC activity also closely tracks autonomic fluctuations (Samuels & Szabadi, 2008a, 2008b). Therefore, the LC has an anatomical and functional capacity to distribute interoceptive signals across the brain.

Among the LC’s afferent inputs, the NTS is uniquely positioned to relay viscerosensory information from the body. While the anatomical projections from the NTS to the LC are well established (Aston-Jones et al., 1991; Van Bockstaele et al., 1999), direct evidence that this pathway regulates behavior has not been investigated. Clinically, this circuit is proposed to contribute to the therapeutic efficacy of vagus nerve stimulation (VNS) in treating epilepsy, depression, and anxiety disorders (Furmaga et al., 2011; George et al., 2007, 2008; Goggins et al., 2022; Grimonprez et al., 2015). Despite these observations, a direct test of whether specific NTS to LC input contributes to behavioral regulation remains absent.

To determine whether NTS projections to the LC influence anxiety-like behavior, we used optogenetics to selectively activate NTS axonal terminals in the LC of rats while simultaneously recording cardiorespiratory function via electrocardiography (ECG) and diaphragmatic electromyography (dEMG). Stimulation occurred prior to behavioral testing to assess whether pre-task activation altered subsequent performance. Anxiety-like behavior was assessed using the elevated plus maze (EPM) and the novelty suppressed feeding test (NSFT). To confirm circuit engagement, we quantified c-Fos expression in the NTS and LC. Overall, we demonstrate that stimulation of NTS to LC terminals does in fact increase different aspects of anxiety-like behaviors.

## Results

### Behavioral and Cardiorespiratory Responses in the EPM

#### Anxiety-like Responses during the EPM

To determine if the neuronal projection from the NTS to LC alters anxiety-like behavior we initially stimulated the cell bodies of NTS neurons transduced with (channel rhodopsin 2) ChR2 prior to entering an EPM. In a separate cohort we then optogenetically stimulated just the axons of these neurons in the LC. Prior to EPM entry, rats received optogenetic stimulation in a pre-test stimulation box for 5 min (470 nm, 10 mW/mm², 20 Hz, 10 ms, 15 s on/5 s off; Fig. 1A). Both NTS^tdTomato^ and NTS-LC^tdTomato^ control rat groups exhibited expected behavior upon EPM entry, immediately moving into the closed arm and spending less than <50% of total time in the open arms, supporting the aversive nature of open-arm exposure (Fig. 1B, C). Optogenetic stimulation of either NTS neurons, or their terminals in the LC, did not alter the percentage of time spent in the open arms (%; NTS^tdTomato^: 37 [26– 48] vs. NTS^ChR2^: 26 [15–38], *t*(22) = 1.421, p = 0.1615 unpaired Student’s t-test, Fig. 1B; NTS-LC^tdTomato^: 41 [31–52] vs. NTS-LC^ChR2^: 34 [23–45]; *t*(18) = 2.090, p = 0.051; unpaired Student’s t-test, Fig. 1B). However, activation of the NTS, or its terminals in the LC, did increase total freezing time (s; NTS-LC^tdTomato^: 12 [-5–31] vs. NTS-LC^ChR2^: 65 [17–112]; *t*(18) = 2.325, p = 0.0320 unpaired Student’s t-test, Fig. 1C); NTS^tdTomato^: 11 [6–16 vs. NTS^ChR2^: 24 [13–36]; *t*(22) = 2.161, p = 0.0419 unpaired Student’s t-test, Fig. 1C). Additionally, grooming was elevated in NTS^ChR2^ rats (*t*(22) = 2.107, p = 0.0467 unpaired Student’s t-test, Fig. 1D), while exploratory reaching was selectively reduced for the NTS-LC^ChR2^ group (*t*(18) = 2.259, p = 0.0365 unpaired Student’s t-test; Fig. 2E). To determine whether physiological responses aligned with the increased anxiety-like behavior during the EPM, we then assessed cardiorespiratory responses upon entering the maze, and during the assay.

**Figure 1.**
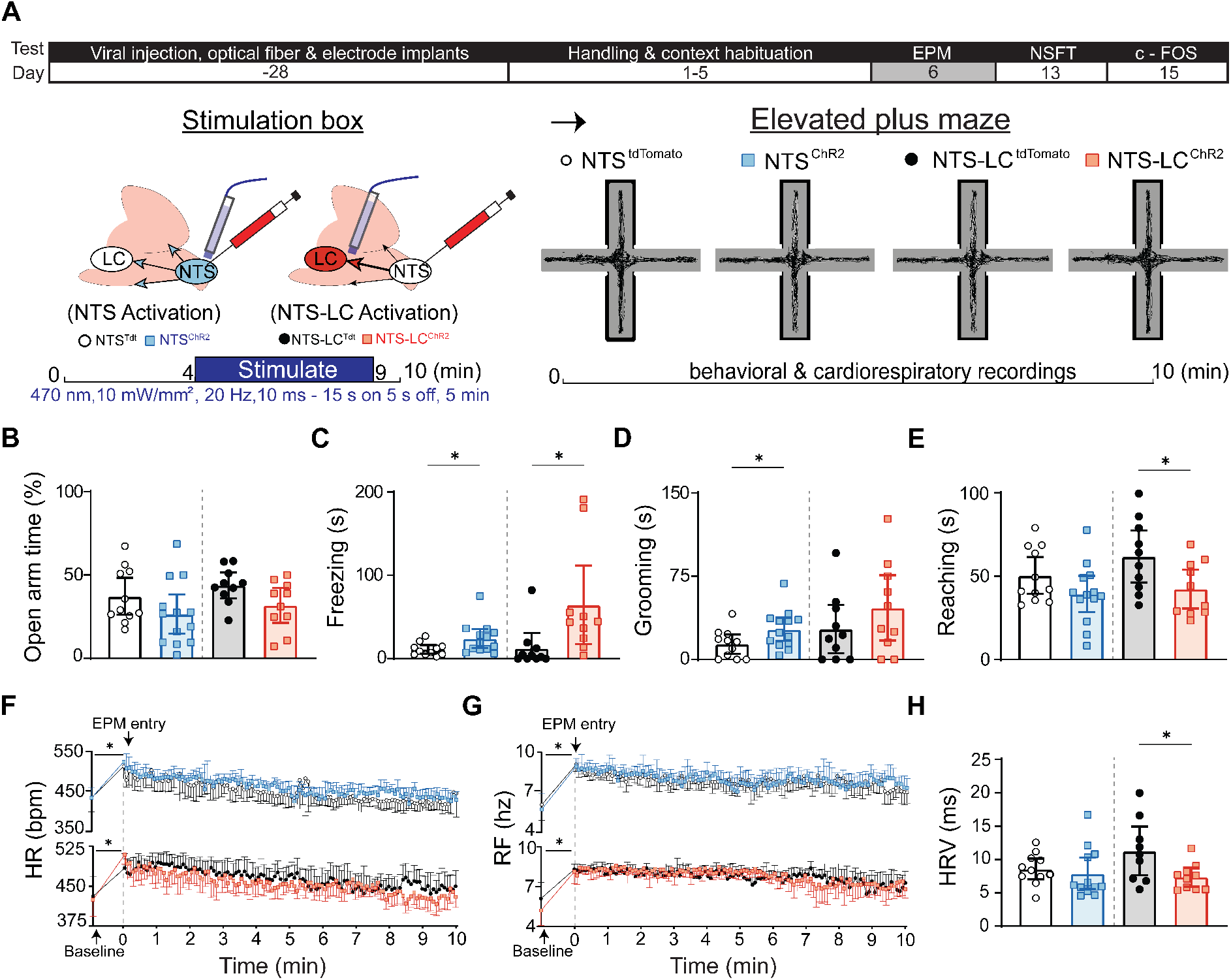
Photoactivation of NTS-LC terminals increases anxiety-like behaviors during the EPM. **(A)** Experimental design, stimulation protocol and representative behavioral traces of NTS^tdTomato^ and NTS^ChR2^ rats during the EPM. Light (470 nm, 10 mW/mm², 20 Hz, 10 ms pulses, 5 min, 15 s on / 5 s off) was delivered to the NTS or NTS-LC terminals before EPM entry. Animals were then placed in the EPM for 10 min. **(B)** Time in open arms (%) for NTS^tdTomato^ (n = 11) vs. NTS^ChR2^ (n = 13) and NTS-LC^tdTomato^ (n = 10) vs. NTS-LC^ChR2^ (n = 10; *: p < 0.05, unpaired Student’s t-test). **(C)** Total freezing (s) during the EPM **for** NTS^tdTomato^ vs. NTS^ChR2^ and NTS-LC^tdTomato^ vs. NTS-LC^ChR2^ (*: p < 0.05, unpaired Student’s t-test). **(D)** Total grooming (s) during the EPM NTS^tdTomato^ vs. NTS^ChR2^ and NTS-LC^tdTomato^ vs. NTS-LC^ChR2^ (*: p < 0.05, unpaired Student’s t-test). **(E)** Total reaching (s) during the EPM for NTS^tdTomato^ vs. NTS^ChR2^ and NTS-LC^tdTomato^ vs. NTS-LC^ChR2^ (*: p < 0.05, unpaired Student’s t-test). **(F-G)** Heart rate (**H**, HR; bpm) and respiratory frequency (**I**, RF; Hz) 15 s before EPM (baseline, in the stimulation box) and throughout the assay for NTS^tdTomato^ (n = 10-11) vs. NTS^ChR2^ (n = 11-12) and NTS-LC^tdTomato^ (n = 10) vs. NTS-LC^ChR2^ (n = 10; *: p < 0.05, two-way RM ANOVA with multiple comparisons, Tukey *post-hoc*).

**Figure 2.**
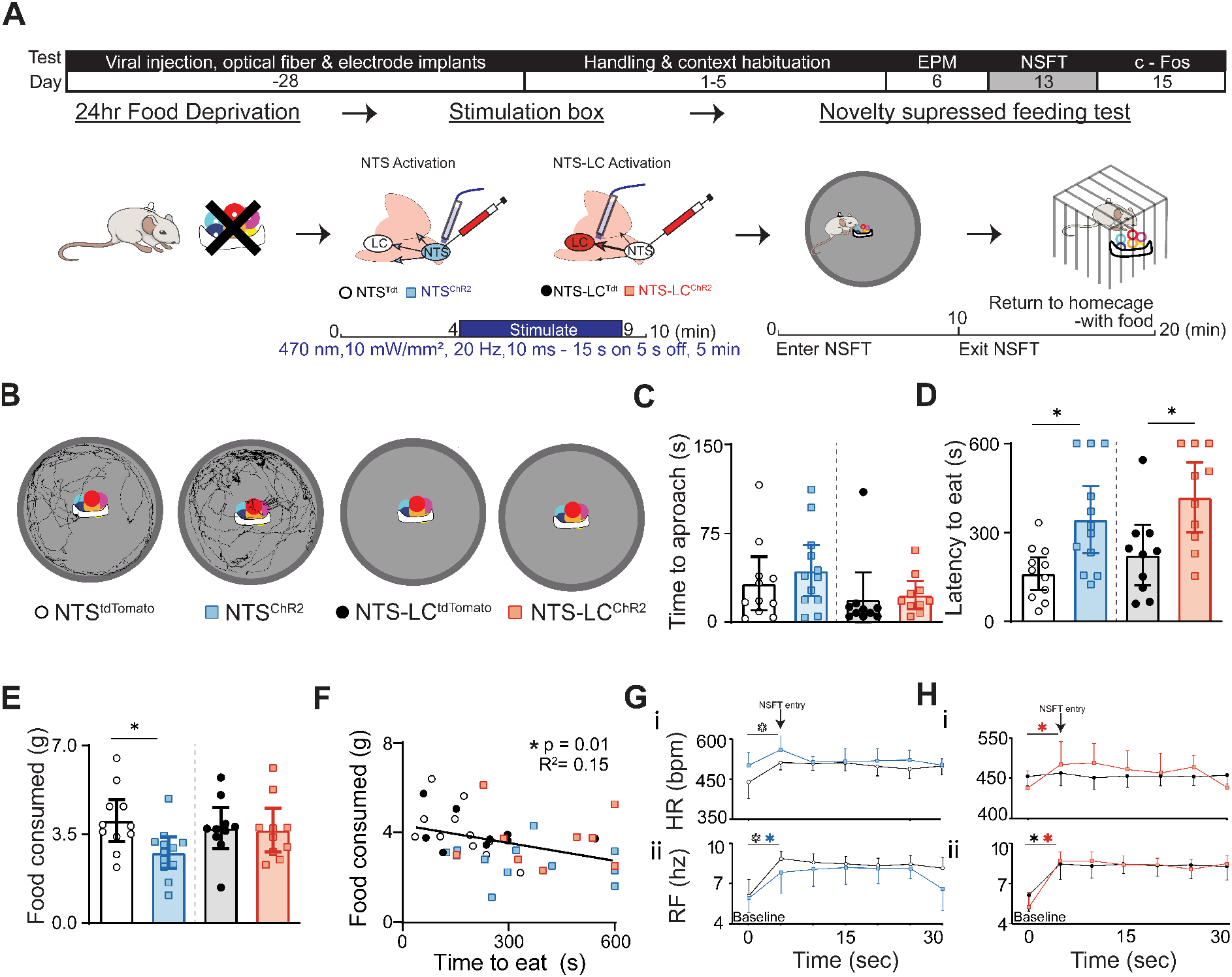
Photoactivation of NTS-LC terminals increases anxiety-like behavior during the NSFT. **(A)** Experimental design and stimulation protocol. **(Ai)** Timeline of data collection. (Aii) Schematic of light administration and NSFT. Animals were food-deprived 24 hours before testing. Light (470 nm, 10 mW/mm², 20 Hz, 10 ms pulses, 5 min, 15 s on / 5 s off) was delivered to the NTS or NTS-LC terminals before NSFT. The NSFT ended when rats ate, or after 600 s. Post-test home cage food consumption was measured for 10 min. **(B)** Representative behavioral traces of NTS^tdTomato^ vs. NTS^ChR2^ and NTS-LC^tdTomato^ vs. NTS-LC^ChR2^ rats during the NSFT. **(C)** Time to approach food in the center for NTS^tdTomato^ (n = 11) vs. NTS^ChR2^ (n = 12) and NTS-LC^tdTomato^ (n = 10) vs. NTS-LC^ChR2^ (n = 10) rats during the NSFT. **(D)** Time to eat in the center of the maze was greater for NTS^ChR2^ (n = 12) and NTS-LC^ChR2^ (n = 10) rats compared to their respective controls (NTS^tdTomato^: n = 11; NTS-LC^tdTomato^: n = 10, *: p < 0.05, unpaired Student’s t-test). **(E)** Post-test home cage feeding was only reduced in NTS^ChR2^ rats (*: p < 0.05, unpaired Student’s t-test). **(F)** Correlation between time to eat and food consumed after the assay (*: p < 0.05, R² = 0.15, Linear regression). **(G-H)** Heart rate (**Gi, Hi;** HR, bpm) and respiratory frequency (**Gii, Hii:** RF, Hz) 15 s before (baseline) and during the first 30 s of the NSFT for **(i)** NTS^tdTomato^ (n = 10) vs. NTS^ChR2^ (n = 12); NTS-LC^tdTomato^ (n = 10) vs NTS-LC^ChR2^ (n = 10) and **(ii)** NTS^tdTomato^ (10) vs. NTS^ChR2^ (n = 11); NTS-LC^tdTomato^ (n = 10) vs NTS-LC^ChR2^ (n = 10); (*: p < 0.05, two-way ANOVA with multiple comparisons, Tukey *post-hoc; symbol color represents the group that changed*).

#### Cardiorespiratory Responses during the EPM

When placed into the EPM, all groups showed immediate increases in HR (beats per minute (bpm); NTS^tdTomato^; pre: 433 [398–468], post: 514 [483–545] vs. NTS^ChR2^; pre: 432 [408–457], post: 525 [498–551]; *F*(1, 20) = 35.98, p = 0.0001 two-way RM ANOVA with multiple comparisons, Tukey *post-hoc*; Fig. 1F; NTS-LC^tdTomato^; pre: 434 [398–471], post: 487 [460–514] vs. NTS-LC^ChR2^; pre: 428 [397–459], post: 513 [494–531]; *F*(1, 18) = 34.65, p = 0.0001 two-way RM ANOVA with multiple comparisons, Tukey *post-hoc*; Fig. 1F) and respiratory frequency (Hz; NTS^tdTomato^; pre: 5.89 [4.74–7.04], post: 8.90 [8.39–9.42] vs. NTS^ChR2^; pre: 5.55 [4.36–6.74], post: 8.74 [8.14–9.34] *F*(1, 19) = 49.53, p = 0.0001 two-way RM ANOVA with multiple comparisons, Tukey *post-hoc*; Fig. 1F; NTS-LC^tdTomato^; pre: 6.05 [4.84–7.28], post: 8.42 [7.25–9.57] vs. NTS-LC^ChR2^; pre: 5.16 [4.05–6.26], post: 8.07 [7.17–8.97]; *F*(1, 18) = 60.02, p = 0.0001, two-way RM ANOVA with multiple comparisons, Tukey *post-hoc*; Fig. 1G). Throughout the EPM a reduction in HR(NTS^tdTomato^ vs. NTS^ChR2^; *F*(6.283, 125.7) = 11.52, p = 0.0001, two-way RM ANOVA, Tukey *post-hoc;* Fig. 1F; NTS-LC^tdTomato^ vs. NTS-LC^ChR2^; F(8.689, 165.1) = 7.996, p = 0.0001, two-way RM ANOVA with multiple comparisons, Tukey *post-hoc*; Fig. 1F) and respiration (NTS^tdTomato^ vs. NTS^ChR2^; *F*(12.99, 259.8) = 4.041, p = 0.0001, two-way RM ANOVA, Tukey *post-hoc;* Fig. 1G; NTS-LC^tdTomato^ vs. NTS-LC^ChR2^; F(9.222, 166.0) = 4.534, p = 0.0001, two-way RM ANOVA with multiple comparisons, Tukey *post-hoc*; Fig. 1G) was observed for all groups. However, during the EPM heart rate variability (HRV) was smaller in NTS-LC^ChR2^ rats (*t*(17) = 2.396, p = 0.0283 unpaired Student’s t-test, Fig. 1H). This cardiorespiratory difference observed during the EPM supports that activation of the NTS terminals in the LC caused a different behavioral state during the EPM.

### Behavioral and Cardiorespiratory Responses in the NSFT

#### Anxiety-like Responses during the NSFT

To determine whether activation of the NTS axonal projections to the LC specifically modulates anxiety-like behavior, rather than indirectly affecting exploration through changes in feeding-related circuits, we performed optogenetic stimulation (470 nm, 10 mW/mm², 20 Hz, 10 ms, 15 s on/5 s off; Fig. 2A) on neurons in the NTS or their terminals in the LC for 5 min prior to the NSFT. After placement in the assay, both control groups (NTS^tdTomato^ and NTS-LC^tdTomato^ rats) initially stayed near the periphery, before approaching the food in the center and taking their first bite, which marked the end of the test (Fig. 2B, C, D). In contrast, NTS^ChR2^ (*t*(22) = 0.7813, p = 0.4434; unpaired Student’s t-test; Fig. 2B, C) and NTS-LC^ChR2^ rats (*t*(18) = 0.3319, p = 0.7438; unpaired Student’s t-test; Fig. 2B, C) approached the food as quickly as controls, but they did not eat immediately; instead, they resumed thigmotaxis, increasing the latency for their first bite of the food in the center (s; NTS^tdTomato^: 164 [104–225] vs. NTS^ChR2^: 344 [56.0–303]; *t*(22) = 3.202, p = 0.0041; unpaired Student’s t-test, Fig. 2D; NTS-LC^tdTomato^: 224 [121–326] vs. NTS-LC^ChR2^: 419 [301–537]; *t*(18) = 2.35, p = 0.011; unpaired Student’s t-test, Fig. 2D).

Following removal from the assay, post-test feeding behavior was assessed to determine whether the increased latency to eat reflected altered feeding drive rather than anxiety-like behavior. Notably, food intake in the first 10 min after reintroduction to the homecage did not differ between NTS-LC^ChR2^ and NTS-LC^tdTomato^ rats ((g); NTS-LC^tdTomato^: 3.7 [2.9 – 4.5] vs. NTS-LC^ChR2^: 3.7 [2.8 – 4.5]; *t*(18) = 0.1601, p = 0.8746; unpaired Student’s t-test; Fig. 2E). In contrast, NTS^ChR2^ rats consumed less food than NTS^tdTomato^ controls (g; NTS^tdTomato^: 4.2 [3.3–4.9] vs. NTS^ChR2^: 2.7 [2.2–3.2]; *t*(21) = 2.475, p = 0.0121; unpaired Student’s t-test; Fig. 2E). There was a weak linear correlation across all rats for time to eat during the NSFT vs. post-test food consumption (R^2^ = 0.15, p = 0.0112; linear regression, Fig 2F). Overall, these data suggest that global NTS activation decreases feeding and increases anxiety-like behavior while stimulating their terminals in the LC just increases anxiety-like behavior. We next examined whether these behavioral differences were accompanied by cardiorespiratory changes either to entering, or during the assay.

#### Cardiorespiratory Responses during the NSFT

Placement in the NSFT apparatus caused distinct cardiorespiratory responses in control groups (NTS^tdTomato^, NTS-LC^tdTomato^). The NTS^tdTomato^ rats showed increases in both HR and respiratory frequency (bpm; pre: 440.8 [403–479] vs. post: 489 [471–506], (*F*(1, 20) = 12.22, p = 0.0023, two-way RM ANOVA with multiple comparisons, Tukey *post-hoc*; Fig. 2G; Hz pre: 6.03 [4.79–7.27] vs. post: 8.77 [8.14–9.40], *F*(1, 14) = 30.33, p = 0.0001, two-way RM ANOVA with multiple comparisons, Tukey *post-hoc*; Fig. 2H), whereas only respiratory frequency increased in NTS-LC^tdTomato^ rats (Hz; pre: 6.05 [4.84–7.27] vs. post: 8.41 [7.25– 9.57], *F*(1, 18) = 42.17, p = 0.0001, two-way RM ANOVA with multiple comparisons, Tukey *post-hoc*; Fig. 2H. Comparatively, broad photoactivation of NTS neurons or their LC-projecting terminals altered the cardiorespiratory responses to NSFT placement. Like controls, both stimulated groups showed different cardiorespiratory profiles upon entering the NSFT arena. NTS^ChR2^ rats failed to exhibit the HR increase observed from NTS^tdTomato^ rats, although respiration increased similarly (Hz; pre: 5.85 [4.76–6.94] vs. post: 7.79 [6.25–9.32], *F*(1, 14) = 30.33, p = 0.0001, two-way RM ANOVA with multiple comparisons, Tukey *post-hoc*; Fig. 2H). In contrast, NTS-LC^ChR2^ rats showed increases in both HR (bpm; pre: 456 [423–488] vs. post: 500 [458–543], *F*(1, 18) = 7.026, p = 0.0163, two-way RM ANOVA with multiple comparisons, Tukey *post-hoc*; Fig. 2H) and respiratory rate (Hz; pre: 5.15 [4.04– 6.26] vs. post: 8.63 [7.95–9.32], *F*(1, 18) = 42.17, p = 0.0001, two-way RM ANOVA with multiple comparisons, Tukey *post-hoc*; Fig. 2H). Lastly, during the first 30 s of the NSFT cardiorespiratory responses were similar between all groups. To further investigate the direct effects of optogenetic stimulation on physiological function we next examined cardiorespiratory dynamics during the pre-test stimulation periods.

#### Cardiorespiratory Effects of Activating the NTS to LC Projection

We next examined whether there were immediate effects of the stimulation on cardiorespiratory parameters measured during the pretest stimulation box. A key difference between the groups is that on NSFT testing day the rats were food deprived for 24 h. Across all testing days light delivery to the control groups and broad photoactivation of the NTS had no effects on HR or respiratory frequency (p > 0.05, two-way RM ANOVA; Fig. 3B-I). However, observed that optically stimulating NTS^ChR2^ rats while unconscious increased HR (*F*(1.516, 30.32) =3.692, p = 0.0478, two-way RM ANOVA with multiple comparisons, Tukey *post-hoc*; Sup. Fig. 1B). In contrast, specific photoactivation of the NTS to LC circuit uniquely decreased HR prior to the NSFT (*F*(8.89, 159.9) = 2.239, p = 0.0226, two-way RM ANOVA with multiple comparisons, Tukey *post-hoc;* Fig. 3E) prior to the NSFT. Unlike broad NTS activation, NTS to LC stimulation in unconscious rats did not alter HR ((*F*(3.180, 50.87) = 0.3919, p = 0.7706, two-way RM ANOVA with multiple comparisons, Tukey *post-hoc;* Sup. Fig. 1D). Lastly, there were no cardiorespiratory effects for any group prior to c-Fos testing at the end of the experimental paradigm (day 15).

**Figure 3.**
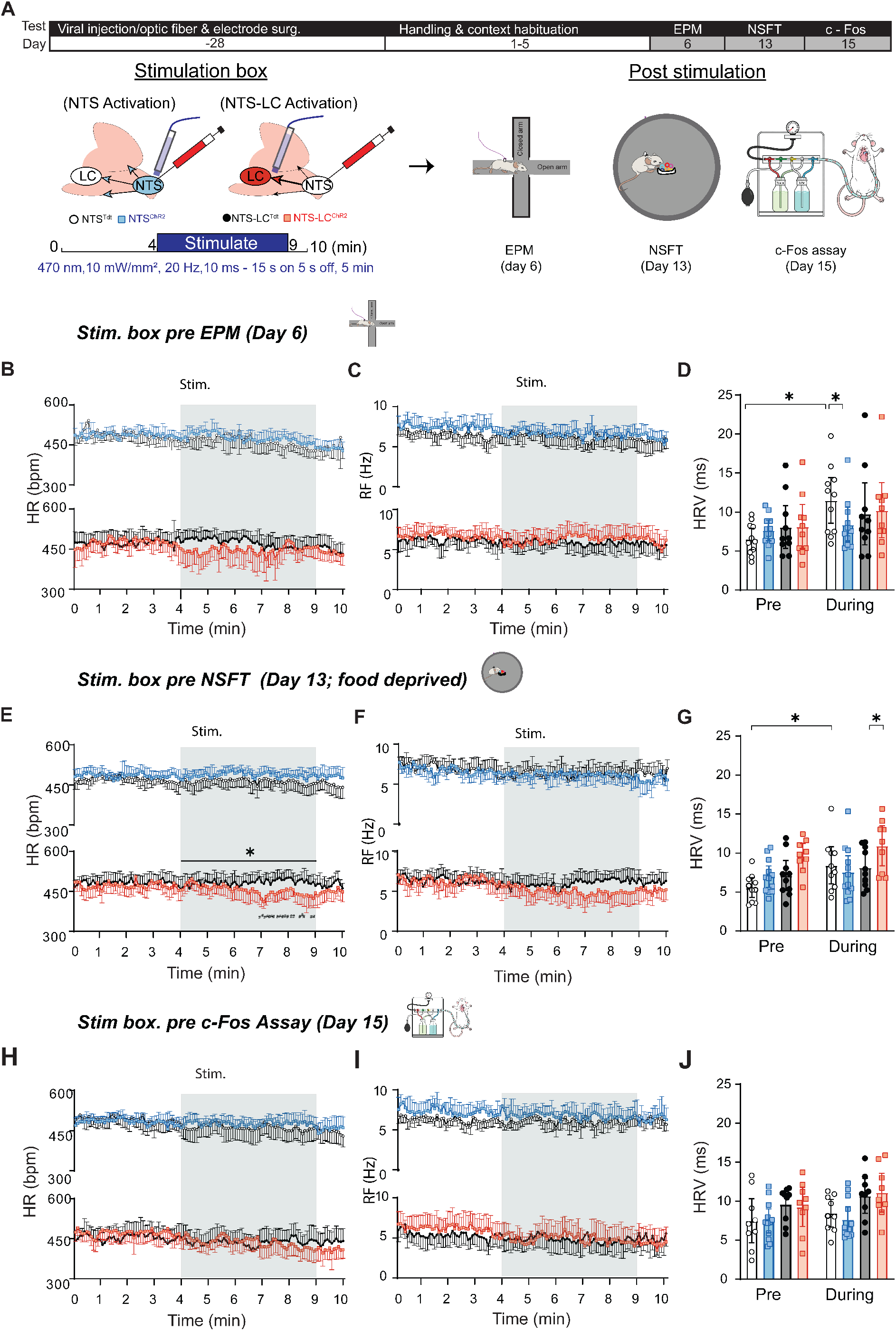
Activation of the NTS-LC pathway has state-dependent cardiovascular effects. (Ai) Experimental timeline, with the shaded box highlighting the period in which the following data were collected. Includes the stimulation protocol on testing days (Day 6: EPM; Day 13: NSFT; Day 15 c-Fos assay), where light was administered to the NTS cell bodies or NTS-LC axonal terminals before behavioral testing. Animals were food-deprived for 24h before NSFT but not EPM or c-Fos assay. Following 5 min in the stimulation box, light (470 nm, 10 mW/mm², 20 Hz, 10 ms pulses, 5 min 15 s on / 5 s off, blue box) was delivered to all rats. After an additional minute, animals were transferred to the behavioral assay. **(B-C; E-F; H-I)** Autonomic responses to optogenetic stimulation, where the grey box in all panels indicates the period when light was delivered to the NTS. **(B)** Heart rate (HR) pre-and during stimulation before the EPM for NTS^tdTomato^ (n = 10) vs. NTS^ChR2^ (n = 12) and NTS-LC^tdTomato^ (n = 10) vs. NTS-LC^ChR2^ (n = 10). **(C)** Respiratory frequency (RF; Hz) pre-and during stimulation before the EPM for NTS^tdTomato^ (n = 11) vs. NTS ^ChR2^ (n = 11) and NTS-LC^tdTomato^ (n = 10) vs. NTS-LC ^ChR2^ (n = 9) **(D)** SDNN pre-and during stimulation before the EPM for NTS^tdTomato^ vs. NTS^ChR2^ and NTS-LC^tdTomato^ vs. NTS-LC^ChR2^ (*: p < 0.05, two-way ANOVA with multiple comparisons, Tukey *post-hoc*). **(E)** HR pre-and during stimulation before the NSFT for NTS^tdTomato^ (n = 10) vs. NTS^ChR2^ (n = 12) and NTS-LC^tdTomato^ (n = 10) vs. NTS-LC^ChR2^ (n = 9; *: p < 0.05 during stimulation between groups; #: depicts p < 0.05 for specific time points during stimulation; two-way ANOVA with multiple comparisons, Tukey *post-hoc*). **(F)** Respiratory frequency across time pre-and during stimulation before the NSFT for NTS^tdTomato^ (n = 8) vs. NTS^ChR2^ (n = 9) and mean respiratory frequency over 10 min in NTS-LC^tdTomato^ (n = 10) vs. NTS-LC^ChR2^ (n = 10). **(G)** SDNN pre-and during stimulation before the NSFT for NTS^tdTomato^ vs. NTS^ChR2^ and NTS-LC^tdTomato^ vs. NTS-LC^ChR2^ (*: p < 0.05 between and within groups; two-way ANOVA with multiple comparisons, Tukey *post-hoc*). **(H)** Heart rate (HR) pre-and during stimulation before the c-Fos assay for NTS^tdTomato^ (n = 9) vs. NTS^ChR2^ (n = 12) and NTS-LC^tdTomato^ (n = 9) vs. NTS-LC^ChR2^ (n = 9). **(I)** Respiratory frequency (RF; Hz) pre-and during stimulation before the c-Fos assay for NTS^tdTomato^ (n = 9) vs. NTS ^ChR2^ (n = 12) and NTS-LC^tdTomato^ (n = 8) vs. NTS-LC ^ChR2^ (n = 9). **(J)** SDNN pre-and during stimulation before the EPM for NTS^tdTomato^ vs. NTS^ChR2^ and NTS-LC^tdTomato^ vs. NTS-LC^ChR2^.

The standard deviation of normal-to-normal intervals (SDNN) was measured before and during stimulation to assess whether optogenetic activation altered heart rate variability (HRV). The effects of photoactivation on HRV differed by group and experimental assay. In NTS^tdTomato^ rats prior to the EPM, HRV increased during stimulation, which was prevented from NTS^ChR2^ rat stimulation (*F*(1, 20) = 6.987, p = 0.0156, two-way RM ANOVA with multiple comparisons, Tukey *post-hoc*; Fig. 3D). In contrast, no significant changes in HRV were observed in any NTS-LC rats during stimulation prior to the EPM. A similar pattern was observed prior to the NSFT for the NTS stimulated group relative to its controls; HRV increased in NTS^tdTomato^ rats (*F*(1, 20) = 5.856, p = 0.0252, two-way RM ANOVA with multiple comparisons, Tukey *post-hoc;* Fig. 3G), but did not differ significantly from NTS^ChR2^ rats However, in this case, NTS-LC^ChR2^ rats showed a significant increase in HRV during stimulation compared to their controls (*F*(1, 17) = 5.785, p = 0.0278, two-way RM ANOVA with multiple comparisons, Tukey *post-hoc;* Fig. 3G). Lastly, on the c-Fos testing day, stimulation had no effect on HRV for either group (p > 0.05, two-way RM ANOVA with multiple comparisons; Fig. 3J). Overall, optogenetic stimulation of the NTS to LC circuit produced distinct cardiorespiratory changes during pre-test stimulation for the EPM and NSFT which appear to be dependent on fed status across groups.

#### Quantifying Neuronal Activation With c-Fos Expression

To confirm neuronal activation following optogenetic stimulation, we assessed c-Fos expression in the NTS and LC (Fig. 4A).

**Figure 4.**
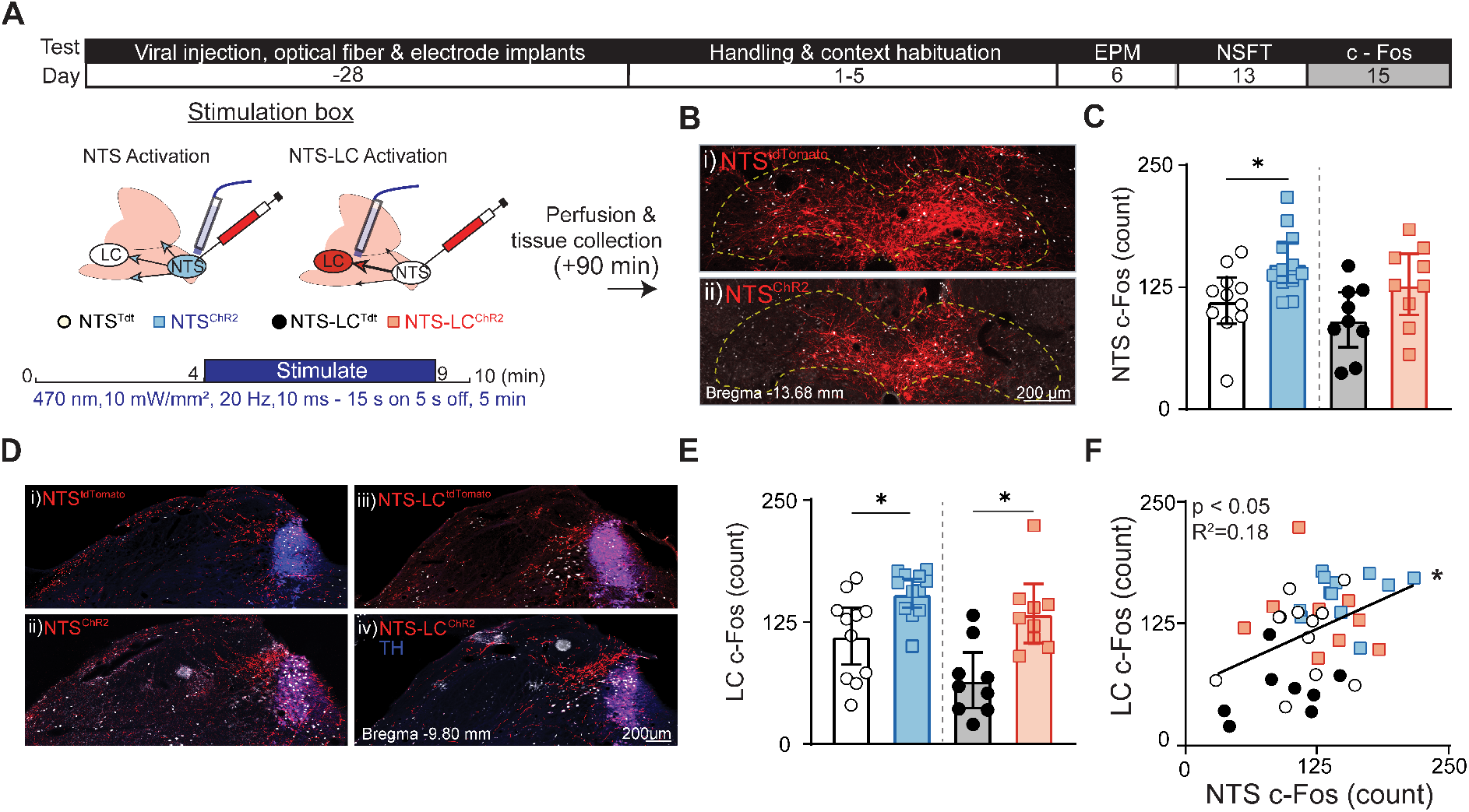
Optogenetic stimulation reliably recruits the NTS-LC circuit. **(A)** Viral strategy and stimulation protocol for c-Fos assay. Includes schematic of viral constructs (AAV2-CAG-ChR2-mCherry for ChR2 activation, AAV2-CBA-tdTomato for controls), transduction timeline (7–8 weeks), and optogenetic stimulation parameters (470 nm, 10 mW/mm², 20 Hz, 10 ms pulses, 5 min, 15 s on / 5 s off). Rats were perfused 90 min after light delivery. **(B)** Representative epifluorescence images of the mid-NTS showing c-Fos-positive neurons (white) colocalized with viral expression (red) in NTS^tdTomato^ vs. NTS^ChR2^ (Bregma:-14.00 mm, dashed line: NTS boundary, scale bar: 200 μm). **(C)** Quantification of c-Fos-positive neurons in the mid-NTS, showing increased expression in NTS^tdTomato^ (n = 11) vs. NTS^ChR2^ (n = 12) and in NTS-LC^tdTomato^ (n = 9) vs. NTS-LC^ChR2^ (n = 9; *p < 0.05, unpaired Student’s t-test). **(D)** Representative epifluorescence images of the LC showing c-Fos-positive neurons (white) with viral expression (red) in NTS^tdTomato^ vs. NTS^ChR2^ and NTS-LC^tdTomato^ vs. NTS-LC^ChR2^ (Bregma:-9.80 mm, scale bar: 200 μm, yellow arrows indicate virally transduced NTS axons near c-Fos-positive cells). **(E)** Quantification of total c-Fos-positive neurons in the LC, showing increased expression in NTS^ChR2^ vs. NTS^tdTomato^ and in NTS-LC^ChR2^ vs. NTS-LC^tdTomato^ (*p < 0.05, unpaired Student’s t-test). **(F)** Linear regression analysis for mNTS c-Fos expression vs LC c-Fos expression (n = 41, R^2^ = 0.18, p < 0.05, linear regression).

#### NTS Activation

Expression of ChR2 was visualized via immunohistochemical detection of the red fluorophore mCherry in NTS^ChR2^ and NTS-LC^ChR2^ rats, and tdTomato in NTS^tdTomato^ and NTS-LC^tdTomato^ rats. Fluorescence was observed along the rostrocaudal extent of the NTS, with the highest density in the medial portion (Fig. 4B, Sup. Fig. 2). Minimal off-target expression was detected in adjacent structures such as the dorsal motor nucleus of the vagus and the area postrema. The control virus (AAV2-CBA-tdTomato) exhibited slightly greater rostrocaudal spread than AAV2-CAG-ChR2-mCherry, despite comparable injection volumes and viral titers (Sup. Fig. 2A, B). Fiber placements in both the NTS and LC were verified post-perfusion in all animals (Sup. Fig. 3).

We observed c-Fos positive neurons within the medial NTS without subnuclear clustering (Fig. 4B). In NTS^ChR2^ rats, photoactivation increased c-Fos expression in the mid-NTS (Bregma-14.00 mm) compared to NTS^tdTomato^ controls (counts; NTS^tdTomato^: 111 [88–135] vs. NTS^ChR2^: 150 [129–171] neurons, *t*(21) = 2.747, p = 0.0121; unpaired Student’s t-test; Fig. 4C). In contrast, stimulation of NTS axonal terminals projecting to the LC did not affect NTS c-Fos expression (counts; NTS-LC^tdTomato^: 100 [60–140] vs. NTS-LC^ChR2^: 127 [97–159] neurons, *t*(16) = 1.295, p = 0.2148; unpaired Student’s t-test; Fig. 4C).

#### LC Activation

Fluorophore-expressing axons from transduced NTS neurons were visible in the LC of all groups, with some axon terminals forming close appositions with LC neuronal somata (Fig. 4D). LC neurons (Bregma:-9.80 mm) exhibited distinct c-Fos expression patterns in the different groups (Fig. 4D). Both control groups had scattered c-Fos immunoreactivity throughout the LC. Whereas photoactivation of the NTS or its terminals in the LC specifically induced c-Fos expression in the dorsal part of the LC. Previous reports indicate the dorsal LC predominantly receives sensory input (Aston-Jones et al., 1991). This expression pattern was larger for both stimulated groups compared to their respective controls (counts; NTS^tdTomato^: 111 [81–140] neurons vs. NTS^ChR2^: 155 [140–170] neurons; *t*(21) = 3.053, p = 0.0060; unpaired Student’s t-test, NTS-LC^tdTomato^: 65 [37 – 94] neurons vs. NTS-LC^ChR2^: 155 [103 – 164] neurons; *t*(16) = 3.791, p = 0.0002; unpaired Student’s t-test; Fig. 4E). We also observed a weak linear correlation between c-Fos-positive neurons in the LC and mid-NTS across all rats (p = 0.0034, R² = 0.20; Fig. 4F), providing further support for functional recruitment of the NTS to LC circuit.

## Discussion

We observed that activation of NTS neurons, including a projection-defined subset targeting the LC, increases anxiety-like behavior and biases physiological outcomes in a state-dependent manner. We speculate that this pathway is providing a route for viscerosensory information processed in the NTS to engage the LC’s diffuse neuromodulatory network to regulate behavioral expression according to peripheral changes. Consistent with this framework, during the EPM and NSFT selective activation of NTS neurons projecting to the LC increased anxiety-like behavior, whereas broader activation of NTS neurons produced both anxiety-like responses and suppression of post-assay feeding. Furthermore, activating the projections from the NTS to the LC only decreased heart rate if food deprived. Together, these findings support an interoceptive model in which NTS to LC signaling shapes both autonomic and behavioral expression.

Here the broad activation of NTS neurons affected both feeding and anxiety-like behaviors. Whereas selective activation of the NTS to LC circuit dissociated these functions during the NSFT, indicating these behaviors are likely mediated by distinct neural pathways. Prior research supports the view that NTS projections regulating feeding and affective behavior can be anatomically and neurochemically segregated. Specifically, NTS neurons expressing cholecystokinin (NTS^CCK^) suppress feeding, but their role in reward and aversion depends on their projections. NTS^CCK^ projections to the parabrachial nucleus (PBN) drive aversion and appetite suppression (Roman et al., 2016; Roman et al., 2017), whereas NTS^CCK^ neurons projecting to the paraventricular nucleus of the hypothalamus (PVN) are linked to satiety and reward (Roman et al., 2017). In contrast, NTS neurons expressing tyrosine hydroxylase (NTS^TH^) regulate feeding in a projection-specific manner but do not alter affective behavior. Specifically, NTS^TH^ neurons projecting to the PVN promoting satiety (Sayar-Atasoy et al., 2023) and NTS^TH^ projecting to the arcuate nucleus of the hypothalamus stimulating feeding (Aklan et al., 2020). Together, these studies and our data support the view that anxiety-like behavior and feeding information can be conveyed by segregated, projection-defined populations within the NTS.

Activation of the NTS to LC circuit increased anxiety-like behaviors across both assays (EPM, NSFT). During the EPM activating this circuit increased freezing and decreased exploratory reaching behavior. It is noteworthy that the open arm time for NTS-LC^ChR2^ rats tended to decrease, although was not significant (p=0.051). We chose to use total freezing time as an additional measure for anxiety-like behavior because anxious rats are reported to freeze more during the EPM (Walf & Frye, 2007). Our observation that NTS-LC activation increases freezing during the EPM is supported by reports that activating the LC increases freezing during anxiety inducing contexts (Soya et al., 2017, 2020; Basu et al., 2024). Which in part seems to be mediated by LC projections to either the basolateral amygdala (Giustino et al., 2020; Soya et al., 2017) or prefrontal cortex (Basu et al., 2024). We also observed that activating the NTS to LC circuit in our study decreased heart rate variability (HRV) which is reported to be lower in anxious rats (Carnevali et al., 2013). Furthermore, latency to eat during the NSFT is commonly interpreted as an anxiety-related measure (Blasco-Serra et al., 2017) and increased following NTS to LC activation. We are confident this is not a secondary effect of activating NTS feeding neurons because we did not observe a change in posttest re-feeding for these rats. Lastly, across both assays, NTS-LC^ChR2^ rats had different cardiorespiratory responses to assay placement. Where food restriction prior to the NSFT may explain the difference in EPM and NSFT stimulation induced outcomes. Overall, these behavioral and physiological observations support the conclusion that activation of NTS projections to the LC increases the expression of anxiety-like behaviors in aversive contexts.

We observed cardiorespiratory responses to optogenetic stimulation that reflect state-dependent modulation, a feature widely reported for neuromodulatory systems (Bradley et al., 2022; McCormick et al., 2020). Consistent with this, broad activation of transduced NTS neurons at their somata produced no measurable cardiorespiratory changes in conscious animals, which may reflect the simultaneous activation of opposing autonomic circuits. In contrast, broad NTS activation in anesthetized animals increased heart rate, suggesting that autonomic outcomes depend on internal states. Comparatively, targeted activation of NTS projections to the LC elicited bradycardia, but only if the rats were food deprived. Food deprivation also caused an increase of heart rate variability for NTS-LC ^ChR2^ rats that was not observed prior to the EPM or c-Fos testing. Notably, unlike broad NTS activation, stimulation of the NTS to LC pathway in anesthetized animals did not evoke measurable cardiorespiratory changes, further supporting the idea that engagement of this circuit depends on ongoing central and peripheral states. Mechanistically, this state dependence may arise because the subset of LC neurons recruited by NTS input depends on the basal activity of the nucleus, which is shaped by factors including arousal, metabolic state, and autonomic tone. Thus, LC responses to viscerosensory input conveyed by the NTS are not fixed but dynamically modulated by ongoing peripheral and central states. Moreover, the linear correlation between c-Fos expression in the medial NTS and LC supports a functional relationship between these nuclei that persists beyond immediate cardiorespiratory readouts.

The suppression of feeding observed following broad NTS activation is likely mediated by NTS projections distinct from the NTS to LC pathway targeted in this study. Here the optical fiber was positioned above the medial NTS, a region where ∼80% of neurons receive viscerosensory vagal afferent input (Bassi et al., 2022; McDougall et al., 2009). The medial NTS contains multiple neurochemically distinct populations known to regulate feeding. These heterogeneous circuits are all likely co-activated by broad stimulation. These include, but are not limited to, NTS^CCK^ (D’Agostino et al., 2016; Roman et al., 2017), NTS^TH^ (Aklan et al., 2020; Chen et al., 2020; Sayar-Atasoy et al., 2023), and NTS neurons expressing glucagon-like protein-1 (Holt et al., 2019; Brierley et al., 2021), pro-opiomelanocortin (Zhan et al., 2013), or prolactin-releasing peptide (Cheng et al., 2021; Mera et al., 2006). All these populations are known to project densely to the hypothalamus, among other regions. Although some of these populations also send inputs to the PBN (Holt et al., 2022), we observed no detectable c-Fos induction in the PBN following broad or projection specific activation of transduced NTS neurons. This suggests that hypothalamic or other downstream projections are more likely to mediate the feeding suppression observed following broad NTS activation. With our experimental protocol, which likely activates multiple neurochemical populations within the ∼1 mm area around the optical fibers tip, we cannot offer useful speculation regarding the specific neurochemical phenotypes involved. Future studies combining genetic and projection-specific approaches will be necessary to determine which NTS neuronal populations and projections are required for the feeding suppression observed following broad NTS activation.

The finding that the NTS to LC circuit promotes anxiety-like behavior highlights this pathway as a circuit of relevance for neuropsychiatric disorders. These conditions are often marked by autonomic dysregulation; including cardiovascular (Dar et al., 2019; Hamer et al., 2012), respiratory (Laviolette et al., 2014; Jerath et al., 2020), and gastrointestinal dysfunction (Fournier et al., 2020). It has been proposed that dysregulation of the brain-body axis contributes to a pathological feedback loop in which autonomic imbalance and psychiatric symptoms perpetuate one another (Santamaría-García et al., 2024; Bonaz et al., 2021). Modulating viscerosensory input may therefore interrupt this maladaptive cycle and improve cognitive outcomes. Supporting this idea, a large body of evidence shows that VNS, which alters viscerosensory signals reaching the NTS, can enhance cognition and alleviate symptoms of depression and anxiety (George et al., 2007; Dorr & Debonnel, 2006; Ruiz, 2024; Butler et al., 2025). Several studies speculate that the NTS to LC pathway plays a key role in mediating these effects, although the precise circuits and modes of engagement remain unresolved (Dorr et al., 2006; Farrand et al., 2020; Roosevelt et al., 2006; Zhi et al., 2024). While our results show that direct activation of the NTS to LC pathway enhances anxiety-like behavior, rather than decreasing it, alternative approaches such as suppressing the pathways activity, biasing LC toward phasic firing, or engaging opposing circuits to decrease tonic firing could yield decreases in anxiety-like behavior. In prior studies, we found that physiologically tuned VNS reduced LC c-Fos expression and anxiety-like behavior (Butler et al., 2025) and decreases extinction memory (Butler et al., 2021), supporting these possibilities. Thus, the apparent discrepancy between our findings and prior VNS studies speculations may reflect differences in how the NTS-LC circuit is engaged, as direct optogenetic activation likely recruits LC neurons or firing modes distinct from those engaged by VNS. Finally, because NTS to LC activation did not affect post-assay feeding, this pathway may offer a means to modulate emotional states without disrupting appetite regulation.

In this study, we show that NTS neurons projecting to the LC modulates aspects of anxiety-like behavior, enhancing freezing in the EPM and prolonging feeding latency in the NSFT without affecting home-cage re-feeding. This dissociation highlights the anatomical segregation of anxiety-like and feeding circuits within the NTS, suggesting that distinct pathways contribute to different aspects of interoceptive processing. Further investigation into the role of the NTS, particularly its interactions with arousal-related circuits, will improve our understanding of how interoception shapes cognition, perception, and adaptive behavior.

## Materials and Methods

### Animal Ethics and Housing

All procedures utilizing animals were approved by the Animal Ethics Committee of the Florey Institute of Neuroscience and Mental Health (approval number: 21-069). All experiments were conducted in accordance with the National Health and Medical Research Council Code of Practice and the Care and Use of Animals for Experimental Purposes in Australia (Australian Code for the Care and Use of Animals for Scientific Purposes, 2013).

A total of 44 male Sprague-Dawley rats (2–3 months old, weight range: 250–370 g, mean weight: 310 ± 40 g) were used in this study. Following surgery, rats were individually housed in a temperature-controlled room (22 ± 1 °C) with *ad libitum* access to standard chow and water on a 12 h:12 h light/dark cycle (lights on: 07:00; lights off: 19:00).

### Experimental Timeline

The experimental timeline is shown in Figure 1A. Rats underwent surgical procedures, including microinjection of virus into the NTS, followed by a 4-week recovery period to allow for viral transduction. Following recovery, all animals underwent five days of habituation to the testing room, stimulation box and handling before being subjected to behavioral testing. Prior to each behavioral test, optogenetic stimulation was delivered in a pre-test stimulation box for 5 min.

### General Anesthesia Protocol

Rats underwent two separate surgical procedures on different days: one for microinjection of recombinant virus and optical fiber implantation, and a second for implanting recording electrodes. All surgical procedures were performed under sterile conditions. Anesthesia was induced with inhalation of 5% isoflurane in room air in a sealed chamber. Rats were then maintained at a surgical anesthetic plane during surgeries with ketamine and medetomidine (0.5 mg/kg and 250 μg/kg, i.m. Pfizer, New York City, USA). Anesthesia depth was monitored to ensure loss of pedal withdrawal and corneal reflexes. Supplemental ketamine (0.5 mg/kg, i.m.) was administered every 30 min as required. Rats were placed on a heating pad (37 °C) to maintain body temperature, and ophthalmic ointment (Polyvisc, Alcon New Zealand Limited, Auckland, New Zealand) was applied to prevent corneal drying.

### Viral Vectors and Stereotaxic Injections

The viral vectors used in this study were adeno-associated viruses (AAV2) made in our laboratory using published protocols (Ngo et al., 2020; Zolotukhin et al., 1999). For this study, the experimental virus included a ubiquitous cytomegalovirus/chicken ß-actin (CAG) promotor driving expression of H134R-channel rhodopsin 2 – a blue light activated non-selective cation channel – and a red fluorophore, mCherry (AAV2-CAG-ChR2-mCherry-WPRE, titre: 6.63x x10^12^ VP/ml (Biosafety clearance number: 2018/039). The control virus had a cytomegalovirus/chicken ß-actin based promoter (CBA) driving expression of the red fluorophore, tandem dimer tomato (tdTomato); AAV2-CBA-tdTomato, titre: 2.20×10^11^ VP/ml (Biosafety clearance number: 2018/039).

Stereotaxic injections were performed as previously described (Ngo et al., 2020). Rats were secured in a stereotaxic frame (RWD Life Science, China) with the incisor bar set at-15 mm. The surgical site was shaved and disinfected with povidone-iodine and ethanol before making a midline incision to expose the dorsal surface of the brainstem through the atlanto-occipital membrane. Injections were made relative to the position of *calamus scriptorius* (CS) and extended rostrocaudally along the anterior-posterior (AP) axis, with specific coordinates chosen to target the medial NTS. Injections were made-0.2, 0, +0.10, +0.25, and +0.40 mm from CS along the AP axis. At sites rostral to CS, injections transitioned from unilateral to bilateral. The mediolateral injection coordinates were the same as the AP distance from CS. Dorsoventral (DV) coordinates were referenced from the dorsal surface of the medulla. All injections were made at a depth of 0.4 mm ventral to this surface. A total of 50 nL of virus was injected per site, for a cumulative volume of 400 nL per rat. The sites chosen for representation in the Figures of this manuscript were 14.60, 14.30, 14.00, and 13.70 mm caudal to Bregma.

Injections were made using a pulled glass pipette, with a 30 μm outside diameter at the tip, which was connected to a picospritzer (World Precision Intrsuments, USA) to ensure precise viral delivery, in small volumes, at a rate of 50 nL/min. After completing the injection, the pipette was left in place for 3 min before it was slowly retracted in 0.1 mm steps every 30 s to prevent dorsal spread along the injection tract. The incision was closed with 4-0 absorbable sutures (Surgical Specialties, Taunton Somerset, England), and animals received postoperative analgesia (meloxicam, 1 mg/kg, s.c., Boehringer Ingelheim, Ingelheim am Rhein, Germany).

### Optical Fiber Implantation

Immediately after viral injections, optical fibers (200 μm diameter, 10 mm length, NA 0.22, RWD Life Science, China) were implanted for optogenetic stimulation. For the NTS, care was taken to ensure that the fiber would not protrude into the brainstem with head flexion. A hole was drilled in the middle of the interparietal bone, at its caudal junction where it fuses with the occipital bone, using a dental drill (Duratech, GMA-102727-004, Wisconsin, USA) with a tip diameter of 0.8 mm (Drill bits, round tip, Item No. 980-00003-00, Catalog No. 78042, Meisinger, Hansemannstrasse Germany). Care was taken to avoid damaging the underlying dura or blood vessels. Excess blood or bone was washed with saline so that the dura above the surface of the cerebellum could be visualized. A small slit was made in the dura with the tip of a 30 G needle (30G x 1⁄2”, Reg. Bevel, Order No. NN-3013R, Terumo, Laguna, Philippines). The optical fiber was then inserted at an angle of 28.5 ± 3.3° through the cerebellum so that the fiber would penetrate through the dura and sit perpendicular to the surface of the medial obex (∼14.30 mm caudal to bregma). The fibers were placed so the tip was located just dorsal to the surface of the area postrema with the center of the fiber directly dorsal to CS. For stimulation of the LC, optical fibers were implanted bilaterally, with tips just dorsal to the LC. To accurately target the LC, extracellular multi-unit recordings were made using a glass microelectrode (impedance 3-10 MΩ) filled with 0.9% NaCl. The LC neurons were identified by spontaneous firing at 1-2 Hz and increased spiking to pinch applied to the contralateral paw. Once the location of the LC was confirmed, the optical fibers were positioned 100 μm dorsal to the recording site. The location of the LC was generally ∼ 9.70 mm caudal to bregma, and 1.25 mm lateral to the midline.

Once placed, the fibers were secured using three screws secured in the dorsal surface of the skull and application of dental cement (Vertex, Netherlands). After surgery, rats received postoperative analgesia (meloxicam, 1 mg/kg, s.c.) and were monitored for recovery. Prior to implantation, the light intensity at the tip of each optical fiber during laser stimulation was measured using a photometer (Thorlabs, NJ, USA). We aimed to deliver 10 mW/mm² at the fiber tip.

### Electrode Implantation for Physiological Recordings

After 3 weeks recovery, the rats underwent a second surgery to implant ECG and diaphragmatic electromyography (dEMG) electrodes using methods described previously (Butler et al., 2021; Butler et al., 2025). In brief, electrodes were positioned subcutaneously for ECG and sutured to the left costal diaphragm for dEMG. The leads were tunneled subcutaneously and externalized via a dorsal connector.

### Behavioral assays

#### Behavior Recording

For all behavioral assays, the rat’s behavior was recorded and analyzed using Top Scan Suite (Cleaver Systems, Reston, VA), which quantified movement and time spent in specific zones. The experimenter was blind to the experimental conditions during analysis. All testing environments were maintained at 22 ± 1 °C and 50 ± 5% humidity, with minimized noise. Rats underwent habituation to the testing room for 30 min before each test, and all tests were conducted between 09:00-15:00 h to control for circadian influences.

#### Elevated Plus Maze (EPM)

The EPM consisted of four arms (100 cm length, 15 cm width), elevated 100 cm from the ground. Two arms were enclosed (walls: 40 cm height), and two were open and illuminated (60-70 lux). One min after optogenetic stimulation, rats were placed in the center facing a closed arm, and behavior was recorded for 10 min. The percentage of time spent in open arms, distance traveled, and velocity were analyzed.

#### Novelty Suppressed Feeding Test (NSFT)

The NSFT apparatus was a circular arena (100 cm diameter, 50 cm wall height), illuminated at 100 lux. Rats were food-deprived for 24 h prior to testing. A small bowl containing 10 g of sweet cereal (Fruit Loops, Kelloggs Australia) was placed in the center of the field. The time taken to eat after entry to the field was measured as the latency. The test was terminated after 600 s, regardless of whether the rat ate any food. The rat was returned to its home cage and the total amount of food consumed in the first 10 min was also measured.

### Cardiorespiratory recordings

#### Unconscious recordings

Immediately after recording electrode implantation, and whilst the rat remained anesthetized, dEMG and ECG signals were recorded during light delivery to the NTS or LC. Signals were amplified (1 kHz gain), filtered (dEMG: 50-300 Hz, ECG: 0.5-150 Hz), and digitized at 5 kHz using an A-D converter and Spike2 software (Cambridge Electronic Design, UK). The HR and respiratory rate were analyzed in 1 s bins, comparing the activity 10 s prior to stimulation with 10 s during stimulation.

#### Conscious recordings

The ECG and dEMG were recorded during the pre-test stimulation and each behavioral test. Signals were analyzed in 5 s bins using the recording parameters described for unconscious cardiorespiratory data. We calculated the standard deviation of normal-to-normal intervals (SDNN), a widely used time-domain measure of heart rate variability (HRV). SDNN is defined as the standard deviation of successive R–R intervals (i.e., the intervals between consecutive heartbeats). This metric reflects the overall variability of heart rate (HR) over the analyzed period, capturing both short-and long-term fluctuations in autonomic input to the heart. We analyzed the 4 min preceding stimulation and the 5 min during stimulation, as well as the total 10 min period during the EPM. The following formula was applied to the ECG data.

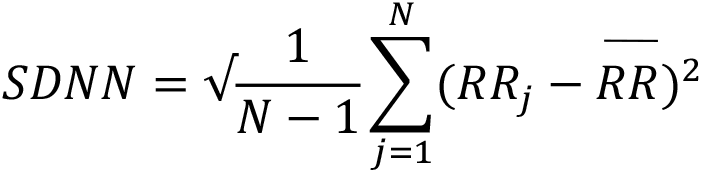

#### Perfusion and Tissue Processing

Ninety min after final optogenetic stimulation, rats were deeply anesthetized with isoflurane and perfused transcardially with 0.1M PBS (300-500 mL), followed by 4% paraformaldehyde in 0.1M PBS w/v (PFA; 300-500 mL). Brains were post-fixed in PFA (4 °C, 24 h), cryoprotected in 30% sucrose (48-72 h), and sectioned coronally (40 μm) using a cryostat (HM525, Microm, Germany). Sections were stored in cryoprotectant solution (30% sucrose, 30% ethylene glycol in 0.1M PBS) until processing.

#### Immunohistochemistry and c-Fos Quantification

Immunohistochemistry was performed as described previously (Butler et al., 2025). Briefly, coronal sections were stained for c-Fos with rabbit anti-c-Fos (1:5000, Synaptic Systems, Göttingen, Germany, catalogue number 226 003, RRID: AB_2231974). The red fluorophore induced by viral expression was amplified with Goat anti-dsRed (1:1000, SICGEN, Coimbra region, Portugal; catalog number AB0040, RRID: AB_2722750) and c-Fos with donkey anti-rabbit Cy5 (1:200, Jackson ImmunoResearch, West Grove, PA, USA; catalog number 711-175-152, RRID: AB_2340607). Incubations were performed on free-floating sections, with an overnight primary antibody incubation at 4 °C, secondary antibody incubation for 4 h at room temperature. Sections were mounted on gelatin-coated glass slides and cover slipped with Cytoseal 60® (ProSciTech, Queensland, Australia). Fluorescence images were acquired using a Zeiss Axioscan 7 slide scanner (Carl Zeiss AG, Oberkochen, Germany). c-Fos-positive cells were quantified using Photoshop (Adobe, CA, USA) with automated thresholding. Regions of interest (NTS or LC) were cropped from coronal sections, and channels were split into red, green and blue (RGB) channels. Thresholding was applied and adjusted to isolate c-Fos-positive nuclei, followed by a median filter to reduce noise. The magic wand tool was then used to select c-Fos-positive nuclei, and automated particle counting was performed for analysis.

#### Experimental Design and Statistical Analysis

Behavioral, physiological, and histological data were analyzed to determine whether optogenetic activation of (1) transduced NTS neurons at the level of their somata (NTS^ChR2^), or (2) NTS axonal terminals within the LC (NTS-LC^ChR2^), produced changes in anxiety-like behavior and/or autonomic output relative to fluorophore-only controls (tdTomato). For all analyses, n refers to the number of rats contributing to the data. For repeated measurements obtained within individual rats (e.g., physiological recordings across time bins, or pre-vs stimulation periods), within-subject factors were treated as repeated measures. All analyses were performed with the experimenter blind to the experimental condition (ChR2 v control groups).

Behavioral outcome measures were freezing behavior, open-arm time, grooming, and exploratory reaching in the EPM and latency to initiate feeding in the NSFT. Seco Physiological outcome measures were HR, respiratory frequency, and heart rate variability quantified as theSDNN. Circuit engagement was assessed by quantification of c-Fos–positive nuclei in the NTS and LC.

Statistical analyses were performed using unpaired Student’s t-tests for planned between-group comparisons, and two-way repeated measures (RM) ANOVA where appropriate for analyses involving within-subject factors, followed by Tukey post hoc tests to account for multiple comparisons. Linear regression was used to examine relationships between behavioral and physiological variables. Statistical significance was defined as p < 0.05. Data are presented as mean ± 95% confidence interval (CI). Final n values for each comparison are reported in the corresponding figure legends.

#### Data Exclusions

Animals were excluded based on predefined criteria, including lack of viral expression, optical fiber misplacement, or excessive noise (>20%) in cardiorespiratory recordings. Final sample sizes for each analysis are reported in the corresponding figure legends. One NTS^ChR2^ rat was excluded from the novelty-suppressed feeding test due to fiber displacement. The number of animals excluded from cardiorespiratory analyses varied between experiments (see figure legends for details).

---

**Supplementary Fig. 1.**
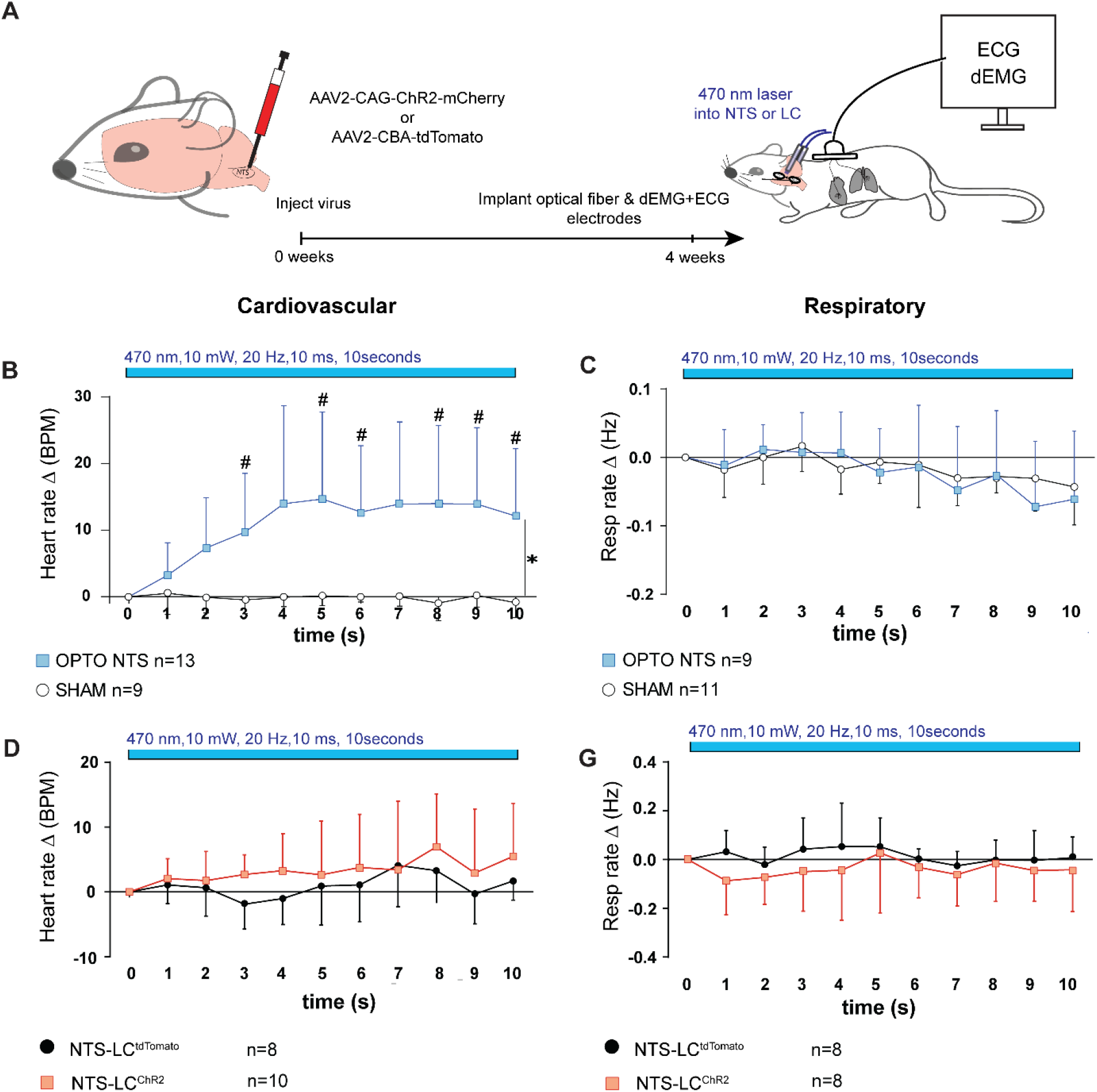
Broad optogenetic activation of NTS increases HR in unconscious rats. **(A)** Schematic of viral injection strategy and experimental setup for ECG and diaphragmatic EMG (dEMG) recordings during optogenetic stimulation. Rats received AAV2-CAG-ChR2-mCherry or control virus (AAV2-CBA-tdTomato) in the NTS, with optical fibers targeting either the NTS or LC terminals. Recordings were conducted 4 weeks post-injection under anesthesia. **(B)** Cardiovascular and (**C**) respiratory responses to 10 s optogenetic stimulation of the NTS (470 nm, 10 mW, 20 Hz, 10 ms pulses) in ChR2 (n = 13) and SHAM (n = 9–11) rats. Broad NTS activation significantly increased heart rate but had no effect on respiratory frequency (#, *: p < 0.05, two-way RM ANOVA with Tukey *post-hoc*). **(D)** Cardiovascular (**E**) and respiratory responses to NTS to LC activation in controls (n = 8–9) and NTS-LC^ChR2^ rats (n = 8–10) rats.

**Supplementary Fig. 2.**
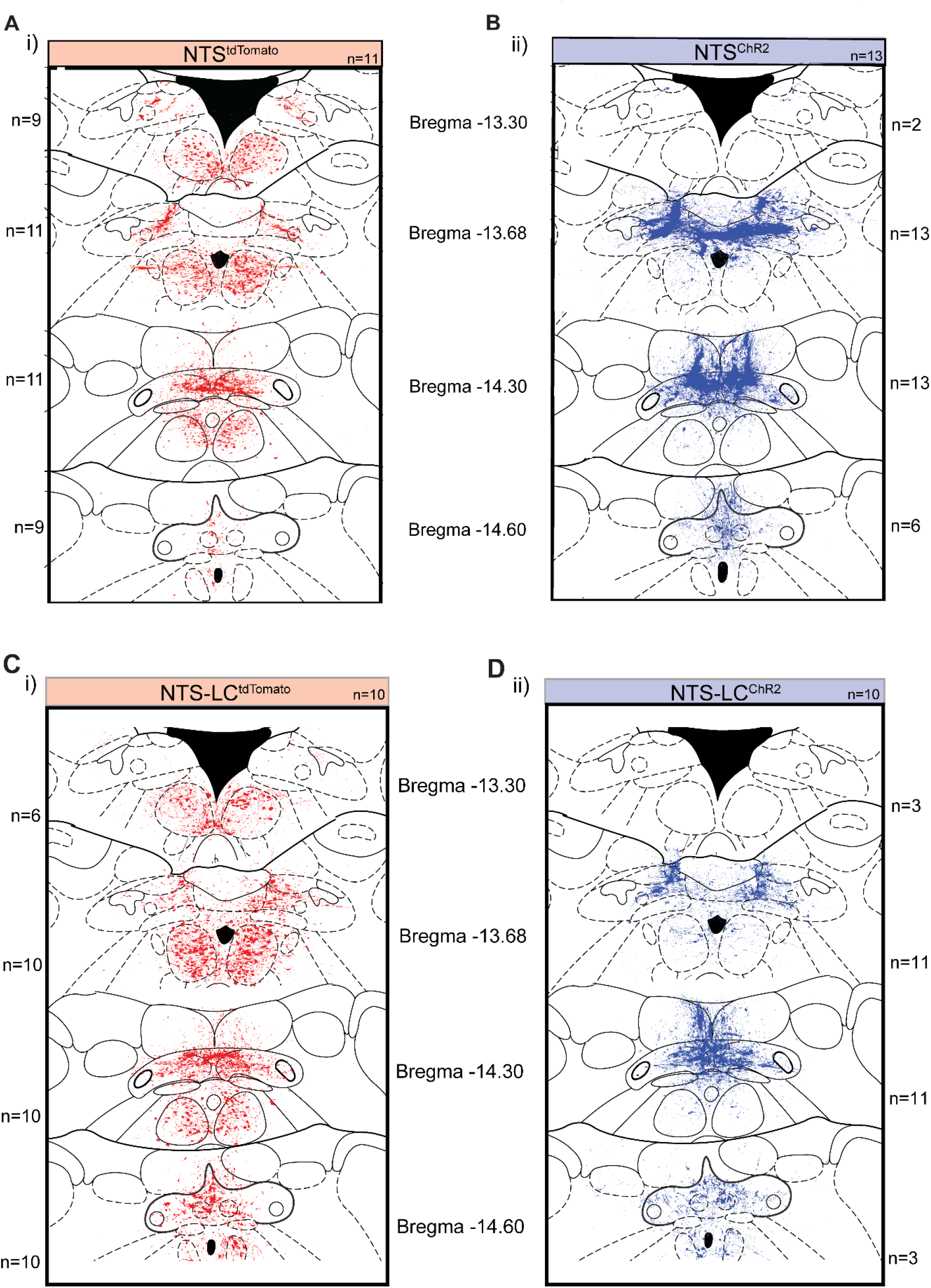
Rostrocaudal distribution of viral expression in the NTS across all experimental groups. (A–B) Viral expression in the NTS for **(A)** tdTomato and **(B)** and ChR2 groups, mapped across Bregma levels: –13.30 mm (n = 9 and 2), –13.68 mm (n = 11 and 13), –14.30 mm (n = 11 and 13), and –14.60 mm (n = 9 and 6), respectively. **(C–D)** Viral expression in **(C)** NTS-LC^tdTomato^ and **(D)** NTS-LC^ChR2^ groups, mapped across Bregma levels: –13.30 mm (n = 10 and 3), –13.68 mm (n = 10 and 11), –14.30 mm (n = 10 and 11), and – 14.60 mm (n = 10 and 3), respectively. Each panel shows overlaid expression from individual animals at each anatomical level, aligned to standard atlas templates to illustrate the consistency and spread of viral transduction across the NTS.

**Supplementary Fig. 3.**
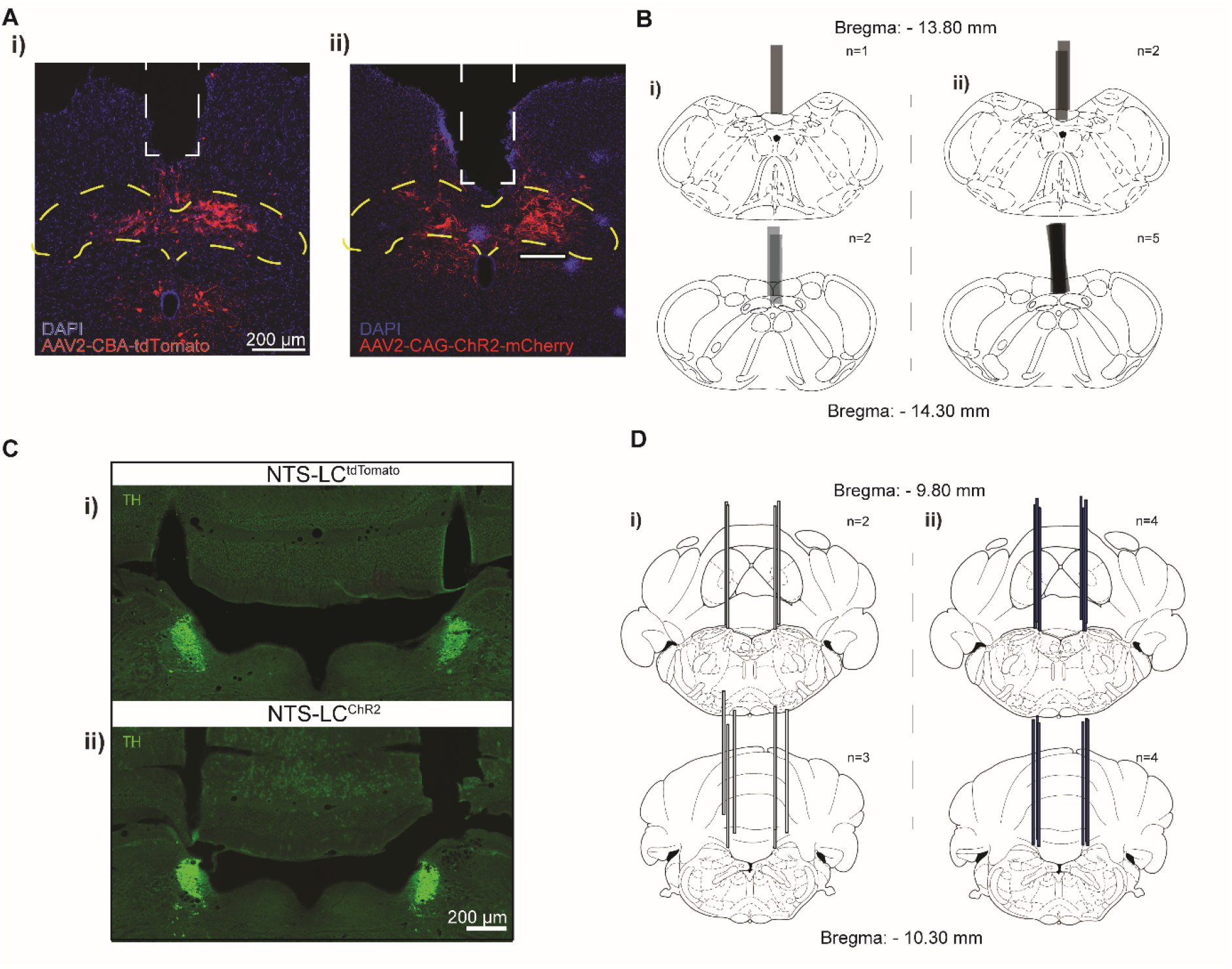
Viral expression and fiber placements for NTS and NTS to LC circuit stimulation. **(A)** Representative epifluorescence images showing viral expression in the NTS for (i) tdTomato control (AAV2-CBA-tdTomato) and (ii) ChR2 (AAV2-CAG-ChR2-mCherry) groups. DAPI (blue) marks nuclei; red indicates viral expression. Dashed yellow lines outline the NTS; white arrows highlight injection spread (scale bar = 200 µm). **(B)** Atlas-mapped optical fiber placements in the NTS for (i) tdTomato and (ii) ChR2 groups across Bregma –13.80 mm (n = 1 and 2) and –14.30 mm (n = 2 and 5), respectively. **(C)** Epifluorescence images of the LC stained for tyrosine hydroxylase (TH, green) confirming projection site targeting and fiber tracts in NTS-LC groups for (i) tdTomato and (ii) ChR2 animals. **(D)** Atlas-mapped optical fiber placements in the LC for (i) tdTomato and (ii) ChR2 groups across Bregma –9.80 mm (n = 2 and 4) and –10.30 mm (n = 3 and 4), respectively.

